# Complete genome sequence of thermophilic cellulolytic actinobacterium *Thermobifida fusca* strain UPMC 901 producing thermostable cellulases

**DOI:** 10.1101/2023.10.16.562643

**Authors:** Mohd Huzairi Mohd Zainudin, Han Ming Gan, Mitsunori Tokura

**Affiliations:** Laboratory of Sustainable Animal Production and Biodiversity, Institute of Tropical Agriculture and Food Security, Universiti Putra Malaysia, 43400, Serdang, Selangor, Malaysia; Patriot Biotech Sdn. Bhd. 47500 Bandar Sunway, Selangor, Malaysia; Research Institute for Bioscience Products & Fine Chemicals, Ajinomoto, 1-1 Suzuki-cho, Kawasaki-ku, Kawasaki-shi 210-8681, Japan

**Author notes:** **Corresponding author:** Mohd Huzairi Mohd Zainudin, Laboratory of Sustainable Animal Production and Biodiversity, Institute of Tropical Agriculture and Food Security, Universiti Putra Malaysia, 43400 UPM Serdang, Selangor, Malaysia.

**Keywords:** Genome sequence, *Thermobifida fusca*, Carbohydrate-active enzymes, Cellulases

## Abstract

Thermophilic cellulolytic bacteria producing thermostable cellulase have multiple potential industrial and biotechnological applications. In this study, we reported the complete genome sequence of the species *Thermobifida fusca* UPMC 901 by using nano-pore sequencing technology. The represented genome is 3.76 Mb with a G + C content of 67.4%. The genome is composed of 1 scaffold with 3,193 protein-coding genes. The results indicated that the genome of the *Thermobifida fusca* gene encodes at least 109 genes for carbohydrate-active enzymes. Of these genes, 38 glycoside hydrolases (GHs), 36 glycosylTransferases (GTs), 21 carbohydrate-binding modules (CBMs), 9 carbohydrate esterases (CEs), and 3 polysaccharide lyases (PL) were identified. In addition, two genes encoding for lytic polysaccharide monooxygenase (LPMO), the copper-dependent enzymes that catalyse oxidative cleavage of recalcitrant crystalline cellulose, that is auxiliary activity (AAs) were also detected in the *T. fusca* genome. This study represented the ninth genome of the species *T. fusca*, and the cellulase genes identified will facilitate the exploration of potential sources from *T. fusca* for industrial and agricultural applications.

## 1. Introduction

Amongst the various cellulolytic microorganisms, thermophilic bacteria have received considerable attention due to their ability to produce thermostable cellulases. Thermostable cellulases have drawn great interest in many commercial applications because of their high specific activity which shortens the hydrolysis period and reduces the risk of contamination and cost of energy for the cooling process after the pre-treatment process. Thus, the exploration of a bacterium harbouring highly thermostable cellulases is extensively carried out by many researchers. Until now several thermophilic cellulolytic bacteria have been isolated from moderate to extreme environmental conditions such as soils, compost heaps, and hot springs (Patel., 2019). The cellulase family was found to play an important role in cellulose biosynthesis as well as biodegradation and multiple applications in various industries such as pulp and paper, food, textile, laundry, and biofuel production.

*Thermobifida fusca*, initially known as *Thermomonospora fusca*, is an aerobic, moderately thermophilic, gram-positive filamentous bacterium. *T. fusca* is a major degrader of the plant cell wall that grows in heated organic material such as compost piles and rotting hay (Wilson, 2004). *T. fusca* secreted multiple types of extracellular cellulases and other carbohydrate-degrading enzymes. These enzyme has been widely studied because of their thermostability, broad pH range, and high activity (Lykidis et al., 2007; Gomez del and Saadeddin, 2014).

*Thermobifida fusca* has been investigated earlier in several studies of lignocellulose degradation including composting and cellulase production. Our previous study of the thermal stability of cellulase demonstrated that crude endoglucanases produced by *T*.*fusca* isolate were highly stable at a prolonged period, indicating the potential of this strain harbouring genes encoding for thermostable enzymes (Zainudin et al., 2019). In addition, it has been suggested that the improved composting of agriculture biomass achieved was enhanced by the presence of several indigenous thermophilic cellulolytic bacteria including *T*.*fusca* (Zainudin et al., 2022). This bacterium has been previously found at the thermophilic phases of the composting process during which the lignocellulose degradation actively took place. Considering all of the information described above, this study aims to analyse the genome sequence of newly isolated *T*.*fusca* UPMC 901, especially on the gene encoding for cellulases to deepen our knowledge and understanding of the role and functions of this bacterium, especially its biomass-degrading enzymes system for lignocellulose degradation.

## 2. Materials and methods

### 2.1 Reviving microbial strain

One ml of Tryptic Soy broth was added into an ampule containing the lyophilized bacteria. The bacterial resuspension was streaked onto two Tryptic Soy agar plates using a sterile inoculating loop and incubated at 50°C for 2 days followed by an additional sub-culture on two new Tryptic Soy agar plates. Bacterial cells were harvested from one plate for genomic DNA extraction, whereas the other plate serves as a backup for glycerol stock preparation.

### 2.2 Genomic DNA extraction

Bacterial cells were harvested from a Tryptic Soy agar plate using a sterile inoculating loop and resuspended in lysis buffer containing 75 mM NaCl, 75 mM Tris-HCL, and 75 mM EDTA followed by the addition of steel and silica beads. The tube containing bacterial cells was subsequently homogenized using a TACO Prep Bead Beater system (GeneReach Biotechnology Corp, Taiwan). The irregular collision of steel and silica beads helps to de-clump the bacterial cells and leads to the complete homogenization of microbial cell walls, respectively. After homogenization, 3% SDS and 200 μg Proteinase K were added to the homogenate and overnight Proteinase K digestion at 55°C was performed. Then, 100 μg of RNAse (PhileKorea Technology, Korea) was added to the bacterial homogenate followed by incubation at 37°C for 15 minutes.

Protein precipitation was performed by the addition of saturated potassium chloride followed by incubation on ice for 15 minutes and centrifugation at 10,000 x g for 15 minutes. Then, chloroform was added to the homogenate to ensure complete protein precipitation, and the homogenate was centrifuged at 10,000 x g for 15 minutes. The supernatant was transferred to a new tube containing isopropanol (1 x vol) for gDNA precipitation. After incubation at room temperature for 10 minutes, the tube was centrifuged at 10,000 x g for 5 minutes. The DNA pellet was washed twice with 75% ethanol, and the DNA pellet was resuspended in 100 μl TE buffer. The gDNA integrity was analysed on a 1X TAE agarose gel.. The DNA was quantified using the DeNovix dsDNA High Sensitivity Kit (Cat# KIT-DSDNA-HIGH-2) and measured on a DeNovix QFX Fluorometer, giving a final concentration of 150 ng/μl.

### 2.3 Size selection

Approximately 10 ug of gDNA was size-selected using 0.15 x vol of SPRI beads in MgCl2-PEG8000 size selection buffer (500 mM MgCl2, 5% PEG-8000) (Stortchevoi et al., 2020). After the removal of the supernatant containing unbound small DNA fragments, the beads were washed with 75% ethanol and DNA was eluted through the addition of 30 μl of TE buffer. The tube was incubated at 37°C for 10 minutes to facilitate the release of DNA from beads into the TE buffer. The final recovered amount of gDNA was 3 μg.

### 2.4 Library preparation and priming of Flongle flow cell

The library was prepared using the Ligation Sequencing Kit (SQK-LSK109). Approximately 1.5 μg of size-selected gDNA was processed using the Nanopore SQK-LSK109 and Native Barcoding Expansion 1-12 kit according to the manufacturer’s instructions with some modifications e.g. the end repair time was increased to 30 minutes at 20oC for and 30 minutes at 65oC. In addition, the ligation time was increased from 10 minutes to 15 minutes. The library was subsequently loaded onto a Flongle flow cell and sequenced for 24 hours.

### 2.5 Data Analysis

Base-calling of the fast5 files used Guppy v4.2.2 (high accuracy mode) without quality filtering. NanoStat v1.4.0 (De Coster et al., 2018) calculated a total of 76,206 reads (315 Mb) with an N50 length of 5,998 bp and a mean read quality of Q11. An initial de novo genome assembly using Flye v2.8.1 (Kolmogorov et al., 2019) (default setting) produced a single contig marked as “circular”. The contig was subsequently polished with Racon v1.4.13 (Vaser et al., 2017), Medaka v.1.2.4 (https://github.com/nanoporetech/medaka), and Homopolish v0.0.1 (Huang et al., 2020). Circularization of the polished genome assembly used a Python script simple_circularise.py (https://github.com/Kzra/Simple-Circularise). The polished genome assembly has a total length of 3,760,927 bp with a GC% of 67.41%. As assessed by BUSCO v4.1.3 (Simão et al., 2015) based on the actinobacteria_odb10 database, the assembled genome is at 98.6% complete.

### 2.6 Identification of carbohydrate-active enzymes (CAZymes)

The carbohydrate-active enzymes in the *T*.*fusca* genome were identified by using the dbCAN2 meta server with Hotpep, HMMER, and DIAMOND search tools.

## 3. Results and Discussion

### 3.1 Characteristic of *T*.*fusca* UPMC 901 genome

In our previous study, several thermophilic cellulolytic bacteria have successfully been isolated during the composting process (Zainudin et al., 2013). Among the isolates, *Thermobifida fusca* UPMC 901, a filamentous, gram-positive actinobacterium was found to exhibit highly thermostable cellulases, particularly endoglucanase (Zainudin et al., 2019). Thus, the genome characteristic of *T. fusca* UPMC 901was carried out in this study to evaluate further their role and functions in lignocellulosic biomass degradation. The final *de novo* genome assembly showed that the *T*.*fusca* genome had a length of 3,760, 927 bp with a G + C content of 67.4%. The completeness of genome assembly and annotation with the BUSCO test suggested a complete annotation set, with 98.6% of the complete BUSCOs and 0.3% fragmented BUSCOs. In total, there were eight *T. fusca* species genomes deposited earlier and published in NCBI genome databases (Table 1). These strains were isolated from cow rumen, composting, and faecal samples of pasture-raised pigs (Martins et al., 2013; Tóth et al., 2013; Stewart et al., 2018). Using nanopore sequencing technology, this study reported the ninth genome of the genus Thermobifida, with the least scaffold. The data indicated that *T. fusca* UPMC 901 had the biggest genome size (3.76 Mb) and highest protein-coding sequences (CDS) of 3,193 in comparison to other *T. fusca* species, indicating the completeness of genome sequence analysis using the nano-pore technique. In addition, based on the NCBI database, we are amongst the first to report the complete genome for assembly level with 100% genome coverage of *T. fusca* species. According to NCBI databases, the *T. fusca* UPMC 901 isolate’s closest neighbours were *Thermobifida fusca* NBRC 14071 (symmetric identity: 96.41%) and *Thermobifida fusca* TM51 (symmetric identity: 96.23%). The genome annotation revealed the presence of 3283 total genes with a total CDS of 3219. Of these genes, 64 RNAs were identified which include 52 tRNAs, 9 rRNAs, and 3 ncRNAs. The assembly also showed 46 pseudogenes and 12 CRISPR arrays (Table 2).

**Table 1.** Genome sequence of *Thermobifida fusca* species. NA = not available

| Organisms name | Organisms group | Strain | Size<br>(Mb) | GC% | Scaffolds | CDS | Assembly<br>level | Accession number |
| --- | --- | --- | --- | --- | --- | --- | --- | --- |
| *Thermobifida fusca* | Bacteria; Terrabacteria; Actinobacteria | JCM 3263 | 3.35 | 66.9 | 1,163 | NA | Contig | BBGT00000000.1 |
| *Thermobifida fusca* | Bacteria; Terrabacteria; Actinobacteria | ZC4RG21 | 3.61 | 67.6 | 204 | 3145 | Contig | QQGVG00000000.1 |
| *Thermobifida fusca* | Bacteria; Terrabacteria; Actinobacteria | RUG14297 | 3.51 | 67.8 | 190 | NA | Contig | CADBTT000000000.1 |
| *Thermobifida fusca* | Bacteria; Terrabacteria; Actinobacteria | SUG1498 | 3.39 | 67.6 | 625 | 3257 | Contig | JAQXTB000000000.1 |
| *Thermobifida fusca* | Bacteria; Terrabacteria; Actinobacteria | TM51 | 3.6 | 67.6 | 88 | 3048 | Contig | AOSG00000000.1 |
| *Thermobifida fusca* | Bacteria; Terrabacteria; Actinobacteria | RUG422 | 3.27 | 67.8 | 453 | 2891 | Contig | ONLN00000000.1 |
| *Thermobifida fusca* | Bacteria; Terrabacteria; Actinobacteria | GAFM-TF-ZOO | 3.64 | 67.5 | 129 | 3080 | Contig | JEMR00000000.1 |
| *Thermobifida fusca* | Bacteria; Terrabacteria; Actinobacteria | NBRC 14071 | 3.64 | 67.5 | 82 | 3106 | Contig | BCWB00000000.1 |
| *Thermobifida fusca* | Bacteria; Terrabacteria; Actinobacteria | UPMC 901 | 3.76 | 67.4 | 1 | 3193 | Complete | CP063232.1 |

**Table 2.** Genome characteristics of *Thermobifida fusca* UPMC 901.

| Items | Value |
| --- | --- |
| Genome size (Mb) | 3.76 |
| GC content (%) | 67.4 |
| Genes (total) | 3283 |
| CDSs (total) | 3219 |
| CDSs (with protein) | 3193 |
| Pseudogene | 46 |
| RNA genes | 64 |
| tRNA | 52 |
| rRNA | 9 |
| nc RNA | 3 |
| CRISPR arrays | 12 |

The carbohydrate-active enzymes (CAZymes) were identified by annotating the genome using dbCAN. Among the 3,193 proteins of *T. fusca* UPMC 901, 109 proteins were annotated as encoding CAZymes of which 39, 36, 21, 9, 3, and 2 proteins belonged to glycoside hydrolyse (GH), glycosyl transferase (GT), Carbohydrate-binding module (CBM), carbohydrate esterase (CE), polysaccharide lyases (PL) and auxiliary activity (AA), respectively (Table 3). These enzymes include 24 families of GH, 9 families of GT, 7 families of CBM, 6 families of CE, 2 families of PL, and 2 families of AA (Fig 1.). Further analysis showed that 37 CAZymes were predicted to be lignocellulose-degrading enzymes, among which cellulases and hemicellulases were found to be the most dominant CAZymes than the other carbohydrate-degrading enzymes. These CAZymes include endoglucanases, β-glucosidases, endo-1,4-β-xylanases, and β-galactosidase. This finding supports the high cellulase and xylanase activity of *T. fusca* UPM 901 as reported earlier by Zainudin et al. (2013, 2019). In addition, enzymes belonging to a family of AA which were referred to as lytic polysaccharide monooxygenase (LPMO) were also detected in the *T. fusca* UPM 901 genome. The comprehensive analysis of proteomics and transcriptomic of cellulolytic aerobic actinobacteria *Streptomyces* SirexAA-E strain isolated from pine-boring woodwasp showed that LPMOs were one of the most abundant proteins detected in the secretomes during the cultivation on pure cellulose and lignocellulose substrates (Takasuka et al., 2013). LPMO is the copper-depend enzyme that catalyze oxidative cleavage of recalcitrant cellulose. The synergistic interaction between LPMO and cellulase is important for hydrolyzing highly crystalline cellulose (Chylenski et al., 2019; Johansen, 2016).

**Table 3.** Carbohydrate hydrolysis enzymes (GH) of *T*.*fusca* UPMC 901 genome.

| GH Family | Product size<br>(No of<br>Amino acid) | Protein / enzymes | Accession<br>number |
| --- | --- | --- | --- |
| GH1 | 484 | Beta – glucosidase | QOS57602.1 |
| GH1 | 463 | Beta – glucosidase | QOS58209.1 |
| GH2 | 830 | Glycoside hydrolase family 2 protein | QOS57581.1 |
| GH3 | 552 | Glycoside hydrolase family 3 C-terminal domain-containing protein | QOS60357.1 |
| GH3,CBM6 | 928 | Glycoside hydrolase family 3 C-terminal domain-containing protein | QOS58188.1 |
| GH4 | 460 | 6-phospho-beta-glucosidase | QOS59251.1 |
| GH5,CBM3 | 616 | Cellulase family glycosylhydrolase | QOS59198.1 |
| GH5,CBM2 | 453 | Cellulase family glycosylhydrolase | QOS57566.1 |
| CBM2,GH5 | 466 | Cellulase family glycosylhydrolase | QOS57567.1 |
| GH6,CBM2 | 441 | Endoglucanase | QOS57731.1 |
| CBM2,GH6 | 596 | Glycoside hydrolase family 6 protein | QOS60187.1 |
| CBM4,GH9,CBM2 | 998 | Glycoside hydrolase family 9 protein | QOS58207.1 |
| GH10 | 399 | Endo-1,4-beta-xylanase | QOS59282.1 |
| GH10,CBM2 | 491 | Endo-1,4-beta-xylanase | QOS59399.1 |
| GH11,CBM2 | 338 | Glycoside hydrolase family 11 protein | QOS57860.1 |
| GH13 | 606 | Maltose alpha-D-glucosyltransferase | QOS60155.1 |
| GH13 | 655 | Alpha-1,4-glucan--maltose-1-phosphate maltosyltransferase | QOS60156.1 |
| GH13 | 544 | Glycoside hydrolase family 13 protein | QOS57506.1 |
| GH13,CBM20 | 605 | Apha amylase C-terminal domain-containing protein | QOS57648.1 |
| CBM48,GH13 | 705 | Glycogen debranching protein GlgX | QOS60572.1 |
| CBM48,GH13 | 724 | 1,4-alpha-glucan branching protein GlgB | QOS60153.1 |
| GH15 | 589 | Glycoside hydrolase family 15 protein | QOS60516.1 |
| GH18 | 451 | Glycoside hydrolase family 18 protein | QOS57538.1 |
| GH18 | 400 | Glycoside hydrolase family 18 protein | QOS60152.1 |
| GH23 | 298 | Lytic transglycosylase domain-containing protein | QOS59081.1 |
| GH23 | 208 | Transglycosylase SLT domain-containing protein | QOS60039.1 |
| GH23 | 386 | Bifunctional lytic transglycosylase/C40 family peptidase | QOS60059.1 |
| GH31 | 765 | Alpha-xylosidase | QOS58193.1 |
| GH42 | 663 | Beta-galactosidase | QOS58195.1 |
| GH43 | 550 | Glycoside hydrolase family 43 protein | QOS58196.1 |
| GH48 | 984 | Cellulose binding domain-containing protein | QOS58492.1 |
| GH63 | 459 | Glycogen debranching protein | QOS58316.1 |
| GH65 | 804 | Glycoside hydrolase family 65 protein | QOS59501.1 |
| GH74,CBM2 | 925 | Cellulose binding domain-containing protein | QOS58192.1 |
| GH77 | 702 | 4-alpha-glucanotransferase | QOS58731.1 |
| GH81 | 758 | Glycoside hydrolase | QOS58662.1 |
| GH95 | 753 | Glycoside hydrolase N-terminal domain-containing protein | QOS58194.1 |
| GH176 | 712 | amylo-alpha-1,6-glucosidase | QOS59445.1 |
| GH178 | 364 | mannan endo-alpha-1,4-mannosidase | QOS60145.1 |

\*GH = Glycoside Hydrolase; CBM: Carbohydrate binding modul

**Fig 1.**
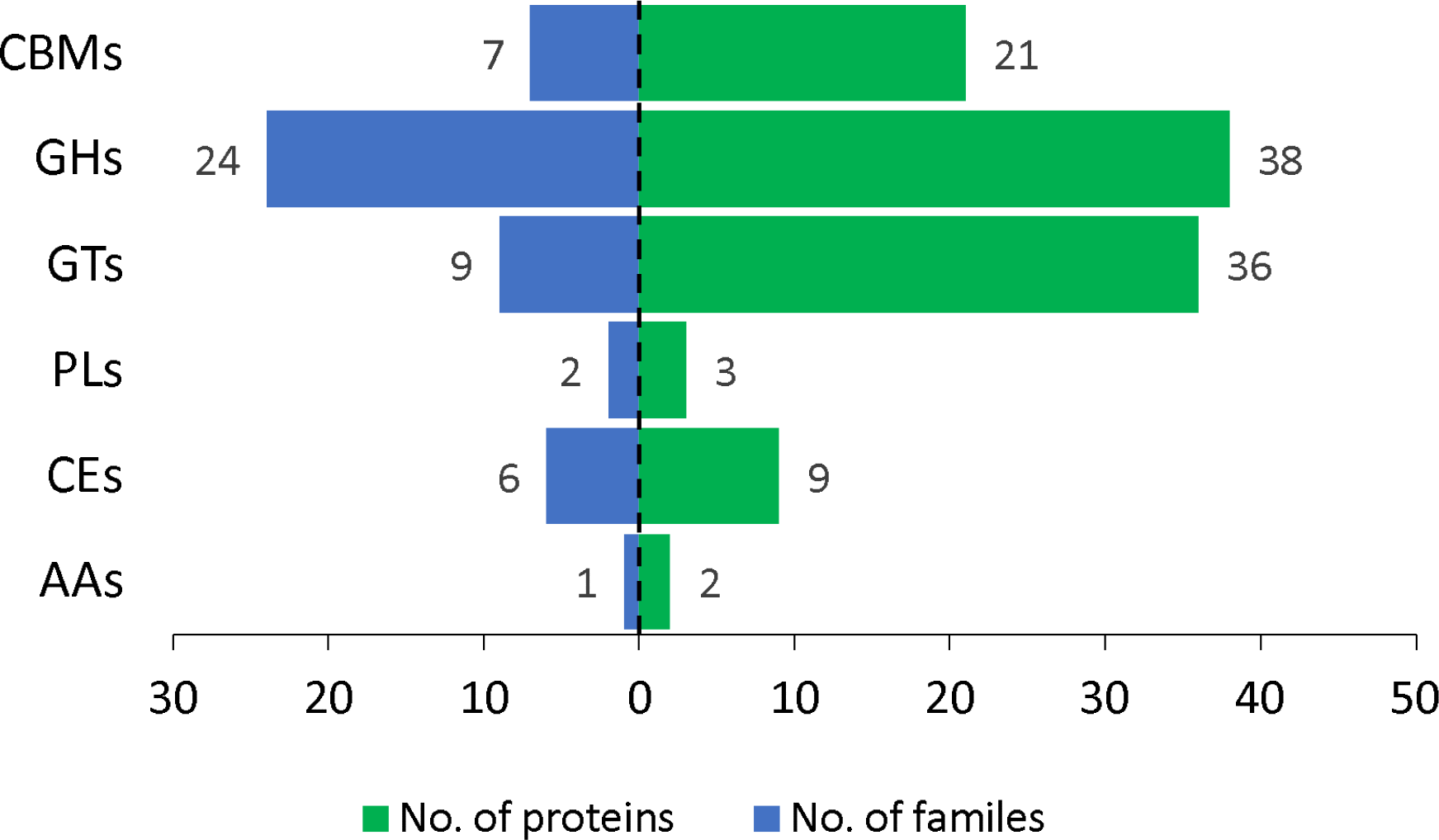
Distribution of carbohydrate-active enzymes (CAZymes) identified in *T. fusca* UPMC 901

In conclusion, the genome information of *T*.*fusca* UPM 901 provides the basis for understanding the properties and functions of this bacterium in assisting the lignocellulosic material degradation. This isolate could be a prospective candidate for remarkable extracellular cellulase and hemicellulose production. Using the nanopore sequencing technology, a complete genome of *T*.*fusca* species was achieved, with fewer contigs and a high rate of BUSCO (98.6%). The genome sequencing revealed 109 gene encoding for CAZymes and their occurrences should be evaluated further independently to ensure the potential of this strain as a feasible candidate for agricultural and industrial application.

## Supporting information

Highlights

## Nucleotide sequence deposition and accession number

The complete genome sequence of *Thermobifida fusca* strain UPMC 901 was submitted and deposited in the NCBI GenBank database under the BioProject PRJNA669597 and BioSample, SAMN16456105 with the accession number CP063232.1.

## CRediT authorship contribution statement

The authors here declare their contributions

Mohd Huzairi Mohd Zainudin: Investigation and writing - original draft and editing. Han Ming Gan: Visualization and Data curation. Mitsunori Tokura: Validation and reviewing.

## Acknowledgment

This project was financially supported by the Universiti Putra Malaysia Grant Putra – IPM (Project number: 9343400). The authors would like to thank the Ministry of Higher Education Malaysia for granting Institute of Tropical Agriculture and Food Security (ITAFoS), UPM as a higher education centre of excellent (HICoE) status.

