## Supplementary material for "Complete genome sequence of thermophilic cellulolytic actinobacterium *Thermobifida fusca* strain UPMC 901 producing thermostable cellulases": Highlights

- *Thermobifida fusca* UPMC 901 as potential strain for cellulase production
- Thermostable cellulase produced by *Thermobifida fusca* UPMC 901
- Genes encoding for carbohydrate-active enzymes (CAZymes)
- Enhancement of lignocellulose degradation
